# Boundary-Sensing Mechanism in Branched Microtubule Networks

**DOI:** 10.1101/2024.08.07.606992

**Authors:** Meisam Zaferani, Ryungeun Song, Ned S. Wingreen, Howard A. Stone, Sabine Petry

**Author notes:** Co-first authors (Meisam Zaferani, Ryungeun Song).

## Abstract

The self-organization of cytoskeletal networks in confined geometries requires sensing and responding to mechanical cues at nanometer to micron scales that allow for dynamic adaptation. Here, we show that the branching of microtubules (MTs) via branching MT nucleation combined with dynamic instability constitutes a boundary-sensing mechanism within confined spaces. Using a nanotechnology platform, we observe the self-organization of a branched MT network in a channel featuring a narrow junction and a closed end. Our observations reveal that branching MT nucleation occurs in the post-narrowing region only if that region exceeds a certain length before it terminates at the channel’s closed end. The length-dependent occurrence of branching MT nucleation arises from the dynamic instability of existing MTs when they interact with the channel’s closed end, combined with the specific timescale required for new MTs to nucleate at a point distant from the closed end, creating a mechanical feedback. Increasing the concentration of the base branching factor TPX2 accelerates nucleation kinetics and thus tunes the minimum length scale needed for occurrence of branching MT nucleation. As such, this feedback not only allows for adaptation to the local geometry, but also allows for tunable formation of MT networks in narrow (micron and submicron scale) channels. However, while a high concentration of TPX2 increases the kinetic rate of branching MT nucleation, it also stabilizes MTs at the channel’s closed end leading to MT growth and nucleation in the reversed direction, and thus hinders boundary sensing. After experimental characterization of boundary-sensing feedback, we propose a minimal model and execute numerical simulations. We investigate how this feedback, wherein growing MTs dynamically sense their physical environment and provide nucleation sites for new MTs, sets a length/time scale that steers the architecture of MT networks in confined spaces. This “search-and-branch” mechanism has implications for the formation of MT networks during neuronal morphogenesis, including axonal growth and the formation of highly branched dendritic networks, as well as for plant development and MT-driven guidance in fungi, and engineering nanotechnologies.

## Introduction

Confinement is known to modulate the self-organization of the cell’s cytoskeleton, which comprises microtubules (MTs), actin, molecular motors, and other associated proteins^1–6^. By rapidly reorganizing such active cytoskeletal networks^7^ a cell can adjust its shape and motility to adapt its function in response to external mechanical cues, e.g. during cell migration and division, and to operate within mechanically complex environments^8–10^. In addition, this adaptive response to mechanical confinement has formed the basis for the engineering of tools to direct the cell’s cytoskeleton and thus its motility^11–14^.

Mechanical sensing under confinement often includes the transformation of a mechanical signal into a biochemical signal^15^. However, cytoskeletal networks can also sense the environment themselves in the absence of biochemical signaling^16^. It has been well-established that the actin cytoskeleton adaptively remodels its architecture in response to physical boundaries^17–22^. MT networks also respond to external mechanical cues^23–28^. For instance, in migrating cells the mechanical properties and stability of individual MTs are altered when exposed to external stress and physical confinement, consequently creating mechanical feedback that enables MT mechanosensation^29–32^. Further, in vitro studies have suggested that centrosomal and motor-driven MT networks, behaving as an active fluid, exhibit adaptive responses to physical boundaries, either directly^33–36^ or through crosstalk with the actin network^37–40^. The acentrosomal networks of MTs in the cortices of plant cells^41^ are also known to self-organize in response to their surrounding confining geometry^42–45^ and to cortical cues such as wall stress^46–48^.

An intriguing aspect of MTs is their ability to autocatalytically self-organize via branching MT nucleation, where new MTs nucleate at shallow angles from existing MTs, and with the same polarity^49–51^. Branching MT nucleation is critical to form the majority of MTs of the mitotic spindle, which segregates chromosomes during cell division^52–54^. Briefly, the GTP-binding protein Ran releases spindle assembly factors around chromosomes, including the branching factors TPX2 and augmin that enable branching MT nucleation in the spindle^55^. In interphase, branching MT nucleation creates cortical MT networks in higher plants^56–58^, makes MTs in the axon^59,60^, which can be meters long, and also creates the highly branched dendritic network in neurons^61,62^. Considering that the self-organization of branched MT networks takes place under cellular confinement, we here investigate whether and how branched MT networks respond to physical boundaries and manage to navigate through confined spaces.

We show that MT nucleation initiated within a certain distance from a physical boundary, coupled with the catastrophe of MTs upon contacting said boundaries, forms a simple feedback system which we call “boundary-sensing.” Our findings are broadly relevant for understanding how MTs drive cellular shape changes and development.

## Results

### Branched MT network formation near a rigid planar boundary

We first quantified the key parameters for comparing MT networks in unconfined and confined spaces. Branched MT networks were initiated in *Xenopus laevis* egg extract by adding 10 µM of constitutively active Ran (RanQ69L). To visualize MTs and their growing plus ends, we added fluorescently labeled tubulin and GFP-end binding protein 1 (EB1) to the extract (Fig. 1A). Using total internal reflection fluorescence (TIRF) microscopy, we observed that branched MT networks started to form within 8-10 minutes after addition of RanQ69L and continued to develop over the period of observation of ∼ 45 min (Fig. 1B).

**Figure 1.**
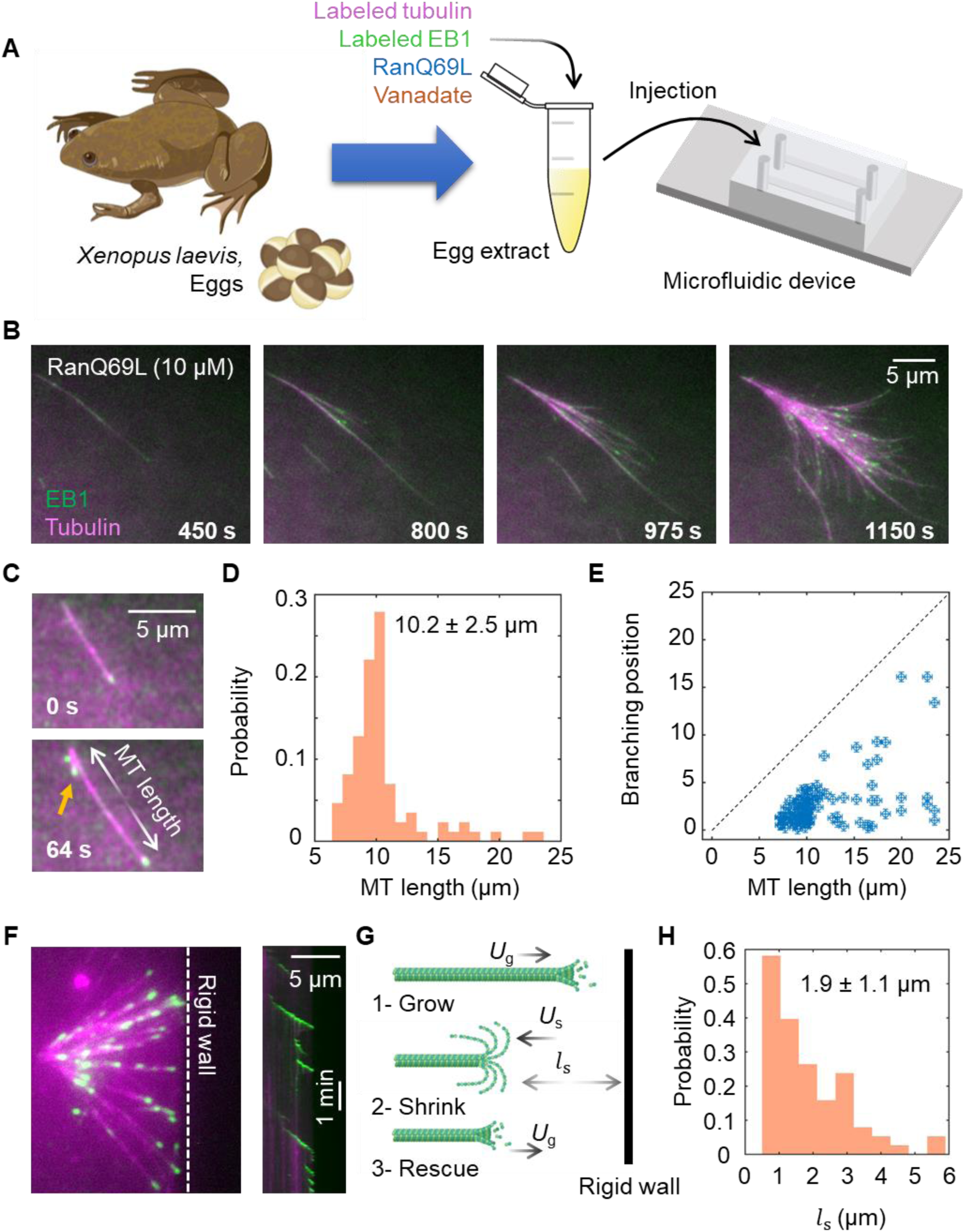
Branched MT network formation near a rigid planar boundary. A) Experimental setup. B) Formation of branched MT networks by addition of 10 mM RanQ69L to the extract system. C) Formation of MT branches occurs on a certain timescale, when mother MTs reach a certain length. Arrow indicates nucleation of daughter MTs. D) Histogram of the mother MT length upon first branch formation (*n* = 37). E) Branching position vs. mother MT’s length (*n* = 37). F) (left) MT network formation toward a rigid wall. (right) Kymograph presents a MT filament approaching the wall with an incidence angle of approximately 90°. G) Illustration of a growing MT shrinking upon contact with a wall and then undergoing rescue. *U*_*g*_ and *U*_*s*_ are growing and shrinking speeds, respectively. *l*_*s*_ is shrinking length. H) Histogram of shrinking length (*n* = 70).

The average length of an existing MT when forming its first branching site was 10.2 ± 2.5 µm (Fig. 1C-D). The positions of branching sites occur close to the minus end of the existing MTs (Fig. 1E), in agreement with our previous measurements – an effect which was shown to be due to a slow on rate of TPX2^63^.

Another key parameter is how MT networks interact with a rigid planar boundary, aka “a wall”. We therefore constructed such a boundary in a simple microfluidic channel and observed branched MT network formation near it. Growing MTs exhibit different behaviors depending on their incidence angles with respect to a wall ^64^. Consistent with our prior work, when incidence angles were acute, MTs were observed to bend and then grow along the wall. When they approached the wall perpendicularly (growth speeds *U*_g_ = 185±27 nm/s, with angle = 87°±5°) they shrank rapidly (*U*_s_ = 0.7-1.5 µm/s) following wall contact (Fig. 1F-G, Movie S1). Then, at some distance from the wall, MTs underwent rescue and regrew until they reached the wall, only to shrink again. We measured the shrinking length (*l*_s_) and found it to be approximately exponentially distributed such that *l*_s_ = 1.9±1.1 µm (Fig. 1H).

### MT network formation in a channel of varying width

We next investigated how both elements, namely branching MT nucleation and MT behavior upon boundary encounter, combine to affect MT networks under confinement, a typical scenario at the cell’s periphery and in its protrusions. To assess this, we confined branched MT networks within a microstructure fabricated at the bottom of a microfluidic chamber (Fig. 2A). The microstructure features a narrow straight channel with one open end, allowing for the entry of a growing MT network into the channel, while the other end is closed, such that MT catastrophe is anticipated upon encounter. The width of the channel is 2 µm, except for a narrow central region with a width of 1 µm (Fig. 2B). By limiting the available space, this narrow region restricts the average number of growing MTs, and then upon expanding again to 2 µm, provides space for branching MT nucleation. We were specifically interested in the number of growing MTs in the post-narrowing region in the presence of the closed end. While this configuration is designed to probe the dynamics of branching MT nucleation under confinement, a channel with varying width and such scale also serves as a generic representation of fine geometrical features found in the neuronal axon and dendrite ^65,66^ and close to the cell wall in plant cells ^57^.

**Figure 2.**
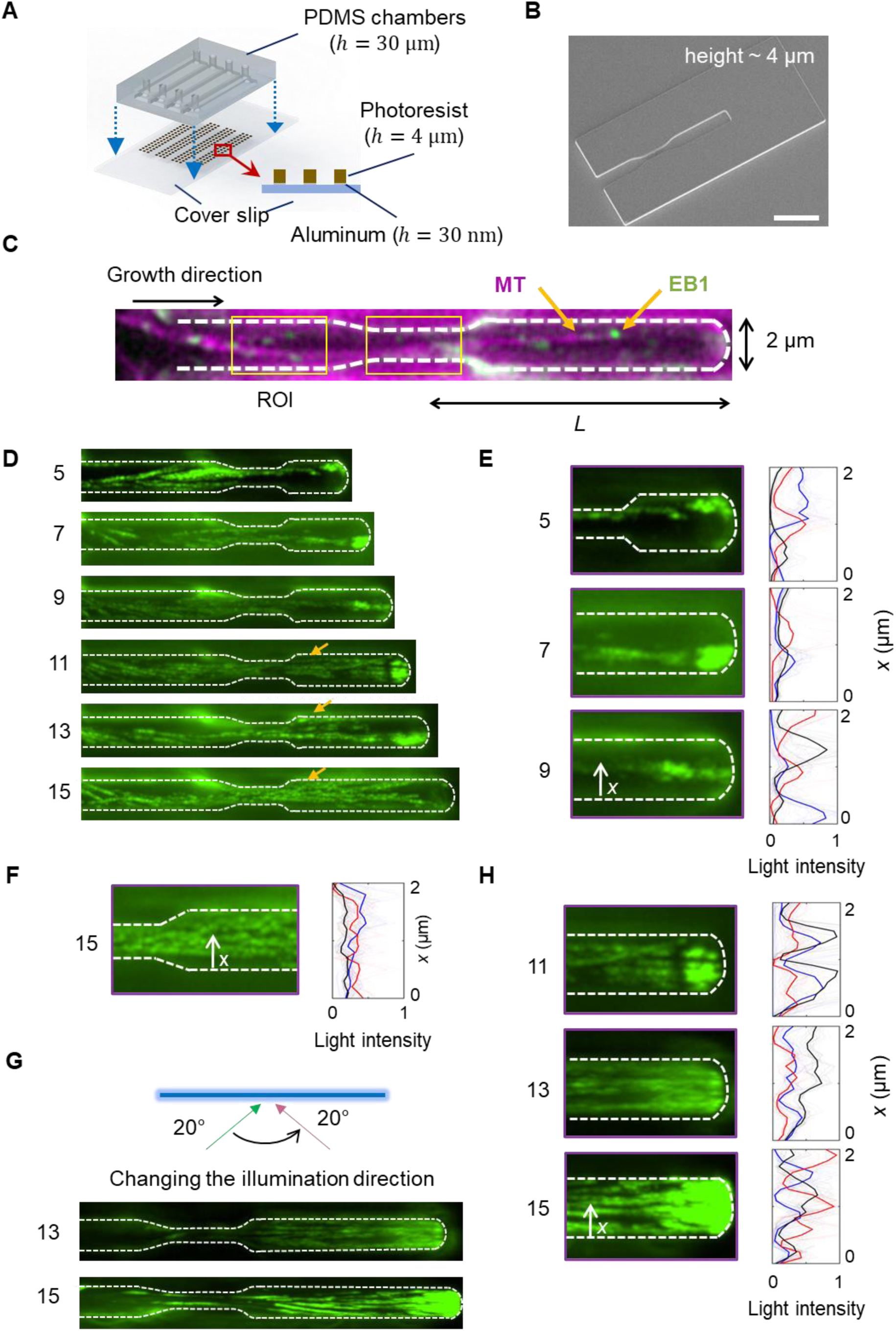
MT network formation in a channel of varying width. A) Schematic of a microfluidic chamber with patterns within, fabricated with photoresist. B) Scanning electron microscopy of a channel with a narrowing region. Scale bar is 5 μm. C) Branched MT network formation inside the channel. Yellow rectangles are regions of interest (ROIs) in pre-narrowing and narrowing regions. D) Self-organization of MT networks after growing through the narrowing region with different post-narrowing region lengths (*L* = 5 to 15 μm). Green lines are overlaid frames taken from EB1 signal (1 fps). Arrows indicate the occurrence of branching MT nucleation in the post-narrowing region when the length of the post-narrowing region exceeds a threshold. Numbers near each image, is the corresponding *L* in µm. E) Light intensity of MT networks at the end of the post-narrowing region for *L* = 5, 7, 9 μm. The light intensity is obtained from the overlaid frames taken from EB1 signal. Different colors represent replicates. F) Light intensity of MT networks at the beginning of the post-narrowing region for *L* = 15 μm. The occurrence of branching MT nucleation in the post-narrowing region results in a wider distribution of growing MTs across the channel width. G) Changing the illumination direction in TIRF, and visualization of the EB1 distribution at the end of the post-narrowing region. H) Light intensity of MT networks at the end of the post-narrowing region for *L* = 11, 13, 15 μm.

The height of the channel was 4 µm, a height which prevents MTs formed on top of the microstructure to enter the channel. We also coated the interface between the coverslip and the fabricated microstructure with a 30 nm thick aluminum layer to suppress the autofluorescence of the microstructures, which enabled imaging of the tubulin signal (Fig. 2C, Movie S2).

The number of growing MTs was inferred by counting the number of EB1 comets, each of which labels a single MT. In the pre-narrowing section, there were 7 ± 2 MTs within a 2 µm × 4 µm region of interest (ROI), while in the narrow section, this number was reduced to 3 ± 1 within a 1 µm × 4 µm ROI (shown in Fig. 2C). We observed that the number of growing MTs in the post-narrowing region depends on the length of this region (*L*), suggesting that branching MT nucleation in the post-narrowing region initiates only if the mother MTs have sufficient room to extend. To investigate this dependency, we fabricated the same channel with various post-narrowing lengths *L*, ranging from 5 to 15 µm (Fig. 2D, Movie S3).

To qualitatively determine the onset of branching MT nucleation in the post-narrowing region as a function of its length *L*, we overlaid frames of EB1 GFP signal at one-minute intervals. For *L* = 5, 7, and 9 µm, no branching MT nucleation was observed (Fig. 2E). Instead, single MTs grew straight from the beginning of the post-narrowing region toward the closed end and remained localized across a small section of the channel width (Fig. 2E). However, for *L* = 11, 13, 15 µm, we observed a wide distribution of EB1 intensity across the channel width at the beginning of the post-narrowing region, indicating the onset of branching MT nucleation at shallow angles (Fig. 2F and arrows in Fig. 2D). Strikingly, this means that increasing the total post-narrowing channel length regulates whether branching MT nucleation takes place at the start of the post-narrowing region.

While we observed EB1 signal at the beginning of the post-narrowing region for *L* = 11, 13, 15 µm, we encountered difficulty observing the entire channel, specifically near the closed end. This problem was mainly due to the high aspect ratio of the microstructure and the corresponding scattered light interfering with the TIRF light path. To address this, we adjusted the illumination direction in the back focal plane of the objective and observed the EB1 signal near the closed end of the channel (Fig. 2G, Movie S4 and S5 for *L* = 13 and 15 µm, respectively). Subsequently, we successfully acquired the EB1 signal at the closed end of the channel, confirming the occurrence of branching MT nucleation and a widespread distribution of the GFP signal across the channel width at the closed end of the channel (Fig. 2H).

To quantify the onset of branching, we measured the number of EB1 comets (*N*_1,2_) in the post-narrowing region for different *L*, at two different regions of interests (ROI_1,2_ = 2 µm × 4 µm). Note that ROI_1,2_ are located at the beginning and near to the closed end of the post-narrowing region, and visualization of EB1 signals was executed at the appropriate illumination direction (Fig. 3A). We observed that at *L* = 11 µm, there is a sharp increase in both *N*_1_ and *N*_2_, which remain elevated at larger *L* values (Fig. 2B, C). This sharp increase of MT numbers at *L* = 11 µm is due to the onset of branching MT nucleation in the post-narrowing region and agrees with the minimal length of the mother MT required for a first branch to form (Fig. 1).

**Figure 3.**
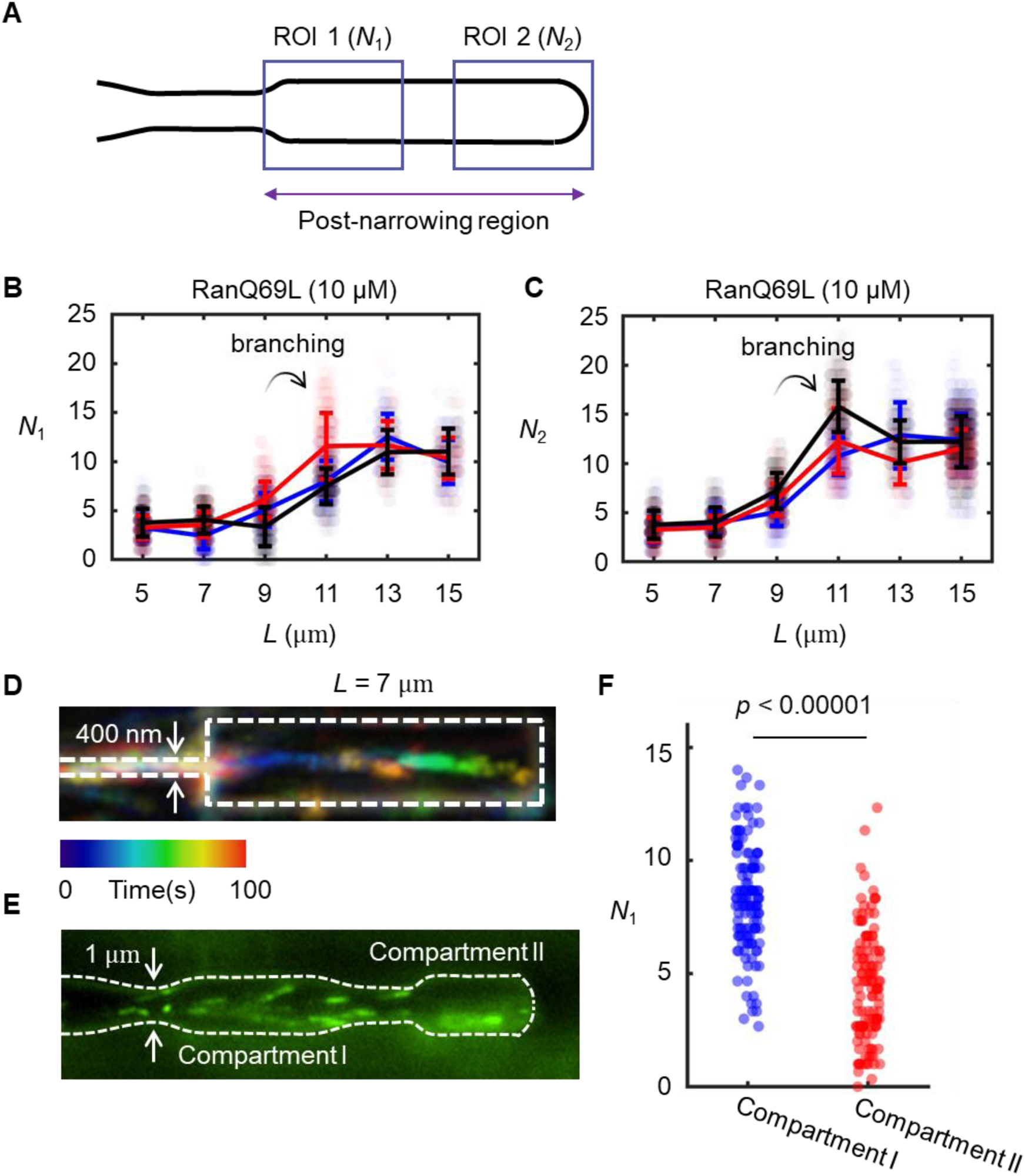
Channel-length-dependent onset of branching MT nucleation in the post-narrowing region. A) Schematic showing two identically sized ROIs within the post-narrowing region. ROI dimensions: 2 µm × 4 µm. B) Number of EB1 spots at the beginning of the post-narrowing region (*N*_1_). C) EB1 spots near the channel end (*N*_2_). Error bars correspond to the standard deviation of *N*_1,2_ and different colors represent replicates. D) MT network growing through a junction of 400 nm width. Typically, few MTs (1-3) were observed growing through the narrowing region, and no branching MT nucleation was observed in the post-narrowing region. E) A single frame showing the EB1 spots in a channel with two narrowing sections. F) Number of EB1 spots within compartments I and II.

To validate that branching MT nucleation does not occur for *L* < 11 µm, we decreased the width of the narrowing region to approximately 400 nm, such that only a very few MTs can pass through this junction. Using a post-narrowing region with *L* = 7 µm, we observed that the number of MTs was indeed reduced to a few (*N* = 1-3) MTs, which did not branch and grew along the center of the channel, where they underwent catastrophe upon encountering the channel’s closed end, if not before (Fig. 3D, Movie S6).

To further assess the influence of the channel’s closed end, we fabricated a channel with two narrowing regions, creating two post-narrowing compartments with lengths of 6.5 and 5 µm, respectively (Fig. 3E, Movie S7). Notably, these two post narrowing compartments differ in that the first one has an open end, whereas the second one has a closed end. Interestingly, we observed that branching MT nucleation events were significantly higher in the first compartment than in the second compartment (Fig. 3F).

These results suggest that branching MT nucleation in the post-narrowing region depends on the distance to the closed end. If this distance is below the minimum length required for branching (*L*_min_ ≈ 10 um), single MTs will encounter the closed end and undergo catastrophe before a branch can form. Thus, branching MT nucleation does not take place in the post-narrowing region. In contrast, if *L* > *L*_min_, MTs take a long enough time to reach the closed end to allow for branching MT nucleation to occur. The initiation of branching MT nucleation compensates for the loss of MTs that have undergone catastrophe at the channel’s closed end. This simple feedback mechanism controls the formation of branched MT networks in confined spaces, enabling an adaptive response to mechanical constrictions. Thus, it serves as a boundary-sensing mechanism.

### The boundary-sensing mechanism is tunable

Extensive evidence suggests that MT stabilizers can regulate the dynamic instability of MTs, which affects how MTs behave at boundaries. However, it is unknown whether tuning the branching MT nucleation rate can also modulate boundary sensing, which we hypothesized to be the case. To test this, we increased the TPX2 concentration in the extract, which leads to a faster formation of TPX2 condensates^67,68^, and thus accelerates recruitment of the branching complex on existing MTs (Fig. 4A).

**Figure 4.**
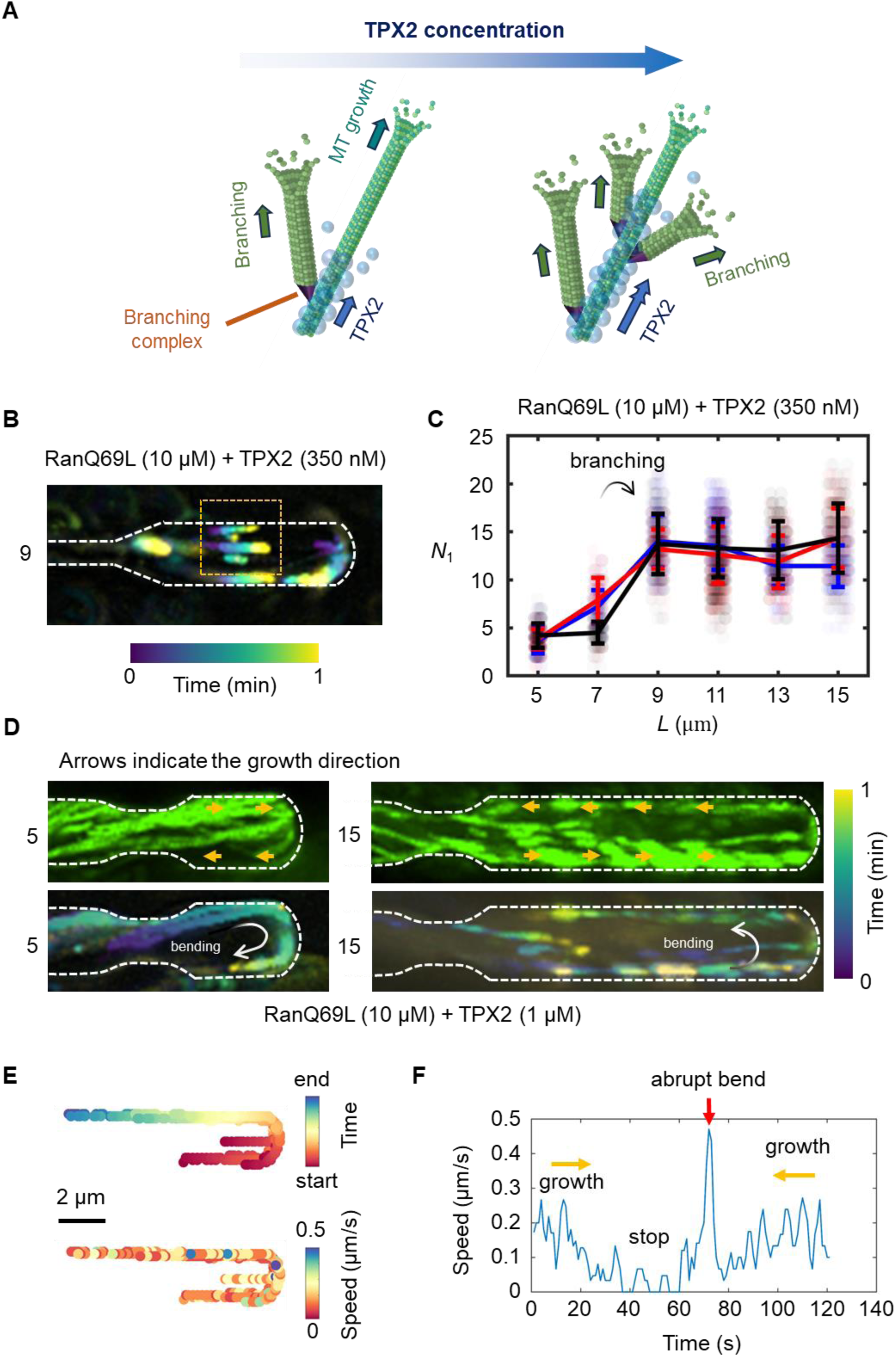
Tuning the adaptive response. A) Schematic of how addition of TPX2 increases the rate of TPX2 layer formation thus faster recruitment of branching complex on existing MTs. B) In the presence of 350 nM TPX2, branching MT nucleation was observed even at *L* = 9 µm. The box highlights nucleation of MT branches, and the color-coded image was obtained by overlaying frames over 1 minute with 1 fps. C) Number of EB1 spots at the beginning of post-narrowing and onset of branching MT nucleation at *L* = 9 µm. Error bars correspond to the standard deviation of *N*_1_ and different colors represent replicates. D) At 1 µM TPX2, branching MT nucleation was observed even at 5 µm, but MTs remain stable after contact with the channel end, eventually resulting in bending and then growth of the MTs in the reverse direction. Arrows indicate the growth direction; the images were obtained by overlaying frames over 1 minute with 1 fps. E) Trajectory of growing MTs, color-coded with respect to time and growth speed. F) The speed profile shown over time corresponds to one of the trajectories depicted in E. The estimated time for the abrupt bend is 15-30 seconds.

Indeed, when we added 350 nM exogenous TPX2, branching MT nucleation could be observed for *L* = 9 µm (Fig. 4B, Movie S8), a post-narrowing channel length for which branching was absent at endogenous TPX2 concentrations (Fig. 2). Next, we counted the number of EB1 comets at the beginning of the post-narrowing region for *L* = 5-15 µm at this elevated TPX2 concentration, which further confirmed a significant shift to *L* = 9 µm required for the onset of branching MT nucleation in the post-narrowing region. Thus, this boundary-sensing mechanism can indeed be tuned to vary the length required for branching to occur.

To determine the extent to which we could shift the length scale for branching MT nucleation to occur, we added 1 µM of TPX2 to the extract. Branching MT nucleation occurred even at *L* = 5 µm (Fig. 4D), confirming the tunable nature of boundary-sensing.

Surprisingly, under these conditions, MTs at the channel ends did not depolymerize but deformed and continued to grow back along the channel in the reverse direction (Fig. 4D, Movie S9). By tracking the EB1 comet for individual, branched MTs (Fig. 4E), we observed that under these conditions, MTs grow and reach the channel end, where their growth pauses for 15-30 seconds. Subsequently, a sharp and rapid bend occurs, ultimately resulting in growth in the reverse direction (Fig. 4F). This high stability of MTs at the boundary can be explained by prior in vitro studies which suggest that high concentrations of TPX2 increase MT lifetime^69^. Therefore, while a unphysiologically high TPX2 concentration enables faster branching, it negatively affects boundary-sensing by stabilizing MTs.

### Theoretical and computational models

Our experimental results show that the combination of dynamic instability near physical boundaries, when coupled with branching MT nucleation at a distant point from the boundary, form a feedback system, which enables MT networks to sense their physical environment in confinement and self-organize accordingly. To quantitatively establish the principles of this feedback, we built a minimal physical model (Fig. 5A), in which a mother MT enters and grows inside a closed channel with a length of *L*, with an average growth speed of *U*_g_. This mother MT shrinks upon contact with the channel’s end with an average shrinkage speed *U*_s_ and a shrinking length *l*_s_ drawn from an exponential probability distribution *p*(*l*_s_), consistent with our experiments (Fig. 1G and H). Then, the expected number of shrinking events required to return to the origin (*x* = 0) is given by:

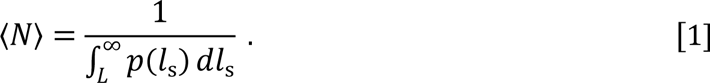

**Figure 5.**
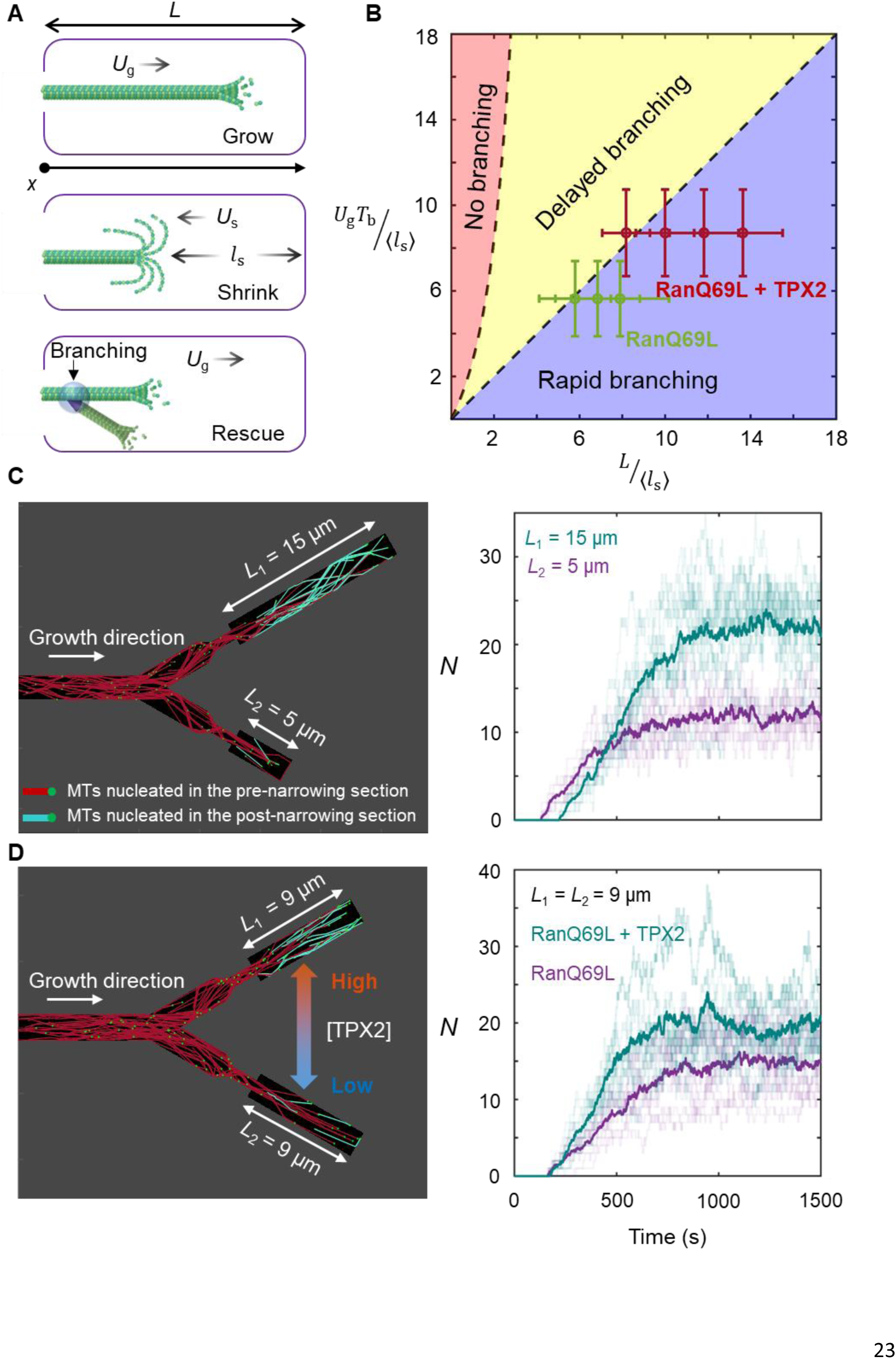
Computational model of MT boundary sensing with experimental comparisons. A) Schematic of our minimal model with B) theoretical phase diagram for branched MT network formation under confinement. Points are experimental measurements for 10 µM RanQ69L and 10 µM RanQ69L + 350 nM TPX2 at different values of *L*. C) Computational simulations of branched MT network formation in a symmetric bifurcation with side channels of different lengths (left panel). The number of MTs (*N*) within 5 µm-wide ROIs near the closed end of each side channel is shown on the right. D) Computational simulations of branched MT network formation in a symmetric bifurcation with higher TPX2 concentration in one side channel (left panel). In the right panels of both C and D, each faint line corresponds to one simulation, while bold lines represent the average of 10 different simulations.

Consequently, the average time needed for the MT to return to the origin can be calculated as:

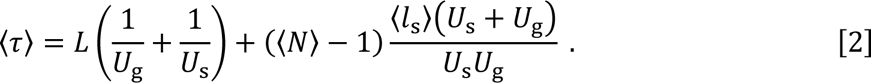

Now, if this 〈τ〉 is substantially shorter than the average branching MT nucleation time (*T*_b_), branching MT nucleation will not occur, because the mother MT will depolymerize before a branch can form. If 〈τ〉 is longer than *T*_b_, branching MT nucleation will generally take place. This condition couples the branching MT nucleation time and confinement length:

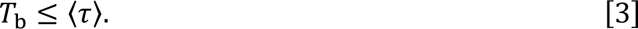

Since *U*_*s*_ ≫ *U*_*g*_, the condition reduces to:

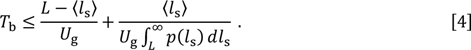

Substituting the exponential distribution for *p*(*l*_*s*_) into the above condition, we obtain the branching MT nucleation condition in a simple form as:

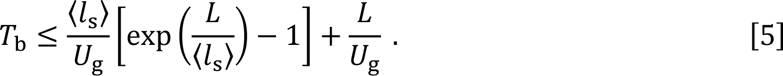

Plotting this curve (Fig. 5B), we appreciate that there is a “no branching” region, where branching MT nucleation may not occur at all.

However, plotting our experimental data, we noticed that while our observations agree with this minimal model, they exist in a sub-regime, where the above condition (Eq. 5) is reduced to: *T*_b_ ≤ *LU*_g_. In this regime, branching MT nucleation occurs before or shortly after the mother MT reaches the end of the channel, which we hence term the “rapid branching regime”. For 〈*τ*〉 ≥ *T*_b_ > *L*/*U*_g_, which we term the “delayed branching regime”, experimental data was not obtained. There might be three reasons for this. First, our assumption that the growth and shrinkage of MTs can persist for extended periods might not be realistic experimentally where photodamage accompanies imaging. Second, it is possible that the time needed to observe the formation of a branched MT network in the delayed branching regime exceeds the duration of our experimental observations, or the branched MT network can only be observed for a short period before it undergoes catastrophe. Third, we used TPX2 to accelerate the branching MT nucleation, but it also impacted stability of MTs. It is possible that using other MT stabilizers that have no impact on the branching MT nucleation reaction, will allow for experimental observations in the delayed branching regime.

Taking advantage of our previously developed simulation framework^64^, we then executed stochastic simulations to assess the consequences of boundary sensing in more complicated geometries. Our first geometry comprises a main channel followed by a symmetric division into two channels each with a width of 2 µm (Movie S10). In a symmetric division, branched MT networks on average divide equally into each side channel. The side channels then feature narrow junctions in the middle, like our experimental setting shown in Fig. 2B, but the side channels have different post-narrowing region lengths: *L* = 5 µm and 15 µm (Fig. 5C). After performing stochastic simulations (ten replicates), we noticed that the MT network only self-organizes in the side channel with the longer post-narrowing region, consistent with our experimental observations. Note that without constricted regions, MT networks would form equally in both channels.

In another scenario, we performed simulations for a completely symmetric division where *L* = 9 µm for both side channels (Fig. 5D, Movie S10). We chose *L* = 9 µm as this length is poised at the critical point for branching. However, we imposed a higher TPX2 concentration in one side channel and found that under this condition, formation of the branched MT network is enhanced in the side channel with highly concentrated TPX2.

## Discussion

In summary, we studied how branched MT networks respond to physical boundaries and navigate through confined spaces. By visualizing the self-organization of branched MT networks in microstructures featuring narrowing junctions and closed ends, we discovered that the onset of branching in the post-narrowing region was determined by the distance to the closed end. The minimum length of the post-narrowing region needed for branching MT nucleation to occur was found to be determined by the MT branching nucleation time and MT catastrophe upon interaction with the wall at the channel’s closed end. Our results suggest that the interplay between dynamic instability, branching MT nucleation, and physical constraints provides a mechanism for boundary-sensing that enables branched MT networks to dynamically adapt to a confined environment.

MT boundary-sensing operates as a feedback loop, enabling MTs to nucleate new branches only if space is available for further growth. This process resembles the “search-and-capture” mechanism previously proposed^70,71^ for individual MTs to dynamically explore and locate targets in their environment. With the addition of branching MT nucleation, this results in a “search- and-branch” mechanism: mother MTs as pioneers grow in confined spaces and when there is enough space, they establish MT networks by providing nucleation sites for daughter MTs. If there is not enough space, the mother MTs simply shrink. We suggest that this search-and-branch mechanism may be relevant for finding and stabilizing nascent neurites before they develop into axons and highly branched dendrites^61,62^.

Another interesting aspect of the search-and-branch mechanism is the force that it can exert when launched. While we did not measure the force generated by growing MT networks, their force can be inferred from the number of growing MTs. Because boundary-sensing can modulate the number and stability of MTs in growing branched networks, it can also determine the force generated by the MT network against the boundary. Specifically, upon the onset of branching MT nucleation under confinement, which results in greater number of MTs (2-3 times), which are overall more spatially and temporally uniform, the force they exert would be consequently both increased and more uniform. Since MT networks coordinate the forces generated at the cellular cortex, such as in neurons^72,73^, filamentous fungi^74^, and higher plants^43^ in response to external mechanical cues, it is conceivable that boundary-sensing via branching MT nucleation contributes to these processes.

Interestingly, the onset of branching MT nucleation could be tuned by adding exogenous TPX2. It is possible that such an effect could be achieved by local translation of TPX2 within cellular extension that are supposed to grow. We also observed that at high TPX2 concentration catastrophes at boundaries no longer occurred, but MTs bent and grew backwards toward their source. This process, at high TPX2 concentrations, results in an array of MTs with mixed polarity, which could be relevant in dendrites which are known to have mixed polarity MT arrays^75,76^.

In the future, it will be exciting to explore in which physiological settings boundary-sensing via branching MT nucleation occurs. It will also be interesting to characterize how boundary-sensing leads to boundary deformation, e.g. of a cell membrane, as has been examined for single MTs^77–79^. In addition, it needs to be investigated how branched MT networks interact with the actin cytoskeleton at cell boundaries, where actin is known to exert force for cell protrusions and locomotion.

## Methods

### (I) Protein Purification and Xenopus Egg Extract

#### Purification of EB1-GFP

To purify EB1-GFP, we employed our previously established methods^80^. Initially, the protein was expressed in *E. coli* (Rosetta 2 strain) at 37°C for 4 hours. Cellular lysis was then achieved using an EmulsiFlex (Avestin) in a lysis buffer composed of 50 mM NaPO4 (pH 7.4), 500 mM NaCl, 20 mM imidazole, 2.5 mM PMSF, 6 mM CME, 1 cOmpleteTM EDTA-free Protease Inhibitor (Sigma), and 1000 U DNAse 1 (Sigma). Subsequent purification involved affinity chromatography with a HisTrap HP 5 mL column (GE Healthcare) using binding buffer containing 50 mM NaPO4 (pH 7.4), 500 mM NaCl, 20 mM imidazole, 2.5 mM PMSF, and 6 mM BME. Elution of the protein was conducted using an elution buffer comprising 50 mM NaPO4 (pH 7.4), 500 mM NaCl, 500 mM imidazole, 2.5 mM PMSF, and 6 mM BME. The resulting peak fractions were combined and loaded onto a Superdex 200 pg 16/600 gel filtration column. Gel filtration was carried out in CSF-XB (10 mM HEPES, pH 7.7, 1 mM MgCl2, 100 mM KCl, 5 mM EGTA) supplemented with 10% (w/v) sucrose.

#### Purification of RanQ69L

RanQ69L, employed for inducing branching MT nucleation, underwent purification following previously reported protocols ^49^. The RanQ69L variant, featuring an N-terminal BFP tag to enhance solubility, was expressed and lysed using the lysis buffer (100 mM tris-HCl, pH 8.0, 450 mM NaCl, 1 mM MgCl2, 1 mM EDTA, 0.5 mM PMSF, 6 mM BME, 200 µM GTP, 1 cOmpleteTM EDTA-free Protease Inhibitor, 1000 U DNAse 1). Affinity purification of the protein was achieved using a StrepTrap HP 5 mL column (GE Healthcare) in binding buffer composed of 100 mM tris-HCl, pH 8.0, 450 mM NaCl, 1 mM MgCl2, 1 mM EDTA, 0.5 mM PMSF, 6 mM BME, and 200 µM GTP. Elution of the bound protein was performed using elution buffer comprising 100 mM tris-HCl, pH 8.0, 450 mM NaCl, 1 mM MgCl2, 1 mM EDTA, 0.5 mM PMSF, 6 mM BME, 200 µM GTP, and 2.5 mM D-desthiobiotin. Finally, the eluted protein underwent overnight dialysis into CSF-XB (10 mM HEPES, pH 7.7, 1 mM MgCl2, 100 mM KCl, 5 mM EGTA) containing 10% (w/v) sucrose.

Tubulin from bovine brain (PurSolutions) was labeled with Alexa 568 NHS ester (GE Healthcare) following previously outlined procedures.

#### Purification of TPX2

To purify TPX2, we employed our previously reported method^67^. The TPX2 construct was cloned as Strep6xHisGFP-TEV-TPX2 fusions at the N-terminus using a modified pST50 vector61 and assembled via Gibson assembly (NEB: E2611L). Transformed into Rosseta2 Escherichia coli cells, the construct was cultured in temperature-controlled incubators at 200 r.p.m. For protein expression, cells were grown at 37 °C (0.5–0.7 OD600), then shifted to 16 °C and induced with 0.75 mM isopropyl-β-D-thiogalactoside for 7 h @ 27 °C. Cell pellets were harvested and promptly frozen for subsequent protein purification. Cell lysis was conducted using an EmulsiFlex (Avestin) in lysis buffer (0.05 M Tris-HCl, 0.015 M Imidazole, 0.75 M NaCl, pH 7.75) supplemented with 0.0002 M phenylmethylsulfonyl fluoride (PMSF), 0.006 M β-mercaptoethanol (βME), cOmplete EDTA-free Protease Inhibitor tablet (Sigma 5056489001), and 1000 U DNase I (Sigma 04716728001). Lysates were centrifuged, and clarified lysate was applied to pre-washed Ni-NTA agarose beads in a gravity column, followed by washing and elution with lysis buffer containing 200 mM Imidazole. Subsequent gel filtration purification was performed using Superdex 200 HiLoad 16/600 columns in CSF-XB buffer. Peak fractions were pooled, concentrated, frozen, and stored at −80 °C.

#### Xenopus egg extract

The branching MT nucleation reactions in the Xenopus egg extract system were conducted according to our previously reported protocols^49^. Briefly, Xenopus egg cytosol, naturally arrested in meiosis II was supplemented with fluorescently labeled tubulin (to a final concentration of 1 µM) for MT visualization. GFP-labeled EB1 (at a final concentration of 100 nM) was introduced to visualize MT growing plus ends. Moreover, RanQ69L (at a final concentration of 10 µM) was included to trigger branching nucleation, while sodium orthovanadate (at a final concentration of 0.5 µM) was included to hinder motors activity and prevent MT gliding on the microfluidic chamber’s bottom surface. The reaction mixture was prepared and kept on ice until it was injected into the microfluidic chamber.

### (II) Microscopy and Image Processing

A Nikon TiE microscope with a 100× 1.49 NA objective was used for Total Internal Reflection Fluorescence (TIRF) microscopy. Images were captured by an Andor Zyla sCMOS camera. The NIS-Elements software (Nikon) was used for image acquisition and capturing images from two channels. To ensure optimal visualization, adjustments were made to brightness, contrast, and acquisition rates (1 and 5 fps) for each experiment. TrackMate software was used to detect, count, and track EB1 comets, with parameters optimized to minimize analysis errors.

Furthermore, the EB1 comet count reported in this work is the number of comets detected in each frame and averaged over 10-second intervals. This averaging reduces the experimental fluctuations associated with comet formation and detection. We also manually checked the results obtained from TrackMate to ensure the validity of the data. The error in the number of EB1 reported by TrackMate was typically ±1, which is below the standard deviation reported for the number of EB1 observed throughout the experiments.

### (III) Microfabrication Process

The microfabrication process follows our previous work^64^, with the addition of an aluminum layer to mitigate autofluorescence from the photoresist structures. Initially, we cleaned the coverslip glasses using Piranha solution and then deposited a 30 nm aluminum layer using a metal sputterer. Subsequently, ∼4 μm thick AZ 4330 photoresist was spin-coated onto the aluminum layer at 4000 rpm for 40 s and soft-baked at 110℃ for 60 s. We used a high-resolution pattern generator (DWL 66+, Heidelberg instruments) to expose the photoresist to the patterned 405 nm UV light. Following this, the photoresist underwent a 5-minute development process in the AZ300MIF developer, while the developer simultaneously etched the aluminum layer under the UV-exposed photoresist. Any residual developer was then rinsed using DI water and dried with nitrogen gas.

The PDMS blocks containing microchannels were fabricated using standard soft-lithography techniques. A 4-inch silicon wafer was spin-coated with SU-8 2025 photoresist at 3000 rpm and then soft-baked on a hot plate at 65℃ for 1 min and 95℃ for 5 min. Subsequently, the SU-8 photoresist was exposed to 375 nm UV light using a medium-resolution pattern generator to generate channel patterns. It was then post-exposure-baked on a hot plate at 65℃ for 1 min and 95℃ for 5 min. before being developed in a SU-8 developer for 5 min. After development, the wafer was rinsed with isopropyl alcohol, dried with nitrogen gas, and then hard-baked on a hot plate at 120℃ for 30 min. PDMS mixture consisting of a 10:1 weight ratio of PDMS base to curing agent was poured onto the SU-8 mold, degassed to remove bubbles, and baked in a convection oven at 70℃ for 5 hours. The cured PDMS was then peeled off from the mold, cut into blocks containing microchannels, and punctured to make inlets and outlets. The PDMS block was bonded with coverslip glass after oxygen plasma treatment.

### (IV) Simulation of MT networks dynamics

To simulate the dynamics of the MT networks within the channel geometry, we modified and utilized the MATLAB software developed in our previous study ^64^. The growth and branching scenarios of MTs were consistent with previous research, but we made a modification to the model for this study: instead of shrinking to a length of 0 when catastrophe occurs, the MTs now shrink by a length *l*_*s*_ sampled from the exponential probability distribution *p*(*l*_s_). Based on our experimental measurements, we set 〈*l*_s_〉 = 2 μm in our simulations.

In simulations of bifurcation geometries with post-narrowing regions of *L =* 5 µm and 15 µm, we initialized the channel’s inlet with MTs, and all parameters related to MT network dynamics were set to the same values as in the previous study ^64^. At each time, the number *N* of MTs in the post-narrowing region was determined by counting the plus-ends of MTs located within a range of 5 µm from the closed end of each branch.

For simulations aimed at observing differences based on TPX2 concentration changes, we increased the binding rate of branching complexes and the nucleation rate by two and √2-fold, respectively, only for MTs entering the upper branch of the channel. This adjustment in branching nucleation rate was intended to evaluate variations based on TPX2 concentration changes. As aforementioned, the number of MTs in the post-narrowing region was determined by counting the plus-ends of MTs located within a range of 5 µm from the closed end of each branch.

